# Spurious correlations in surface-based functional brain imaging

**DOI:** 10.1101/2024.07.09.602799

**Authors:** Jayson Jeganathan, Nikitas C Koussis, Bryan Paton, Sina Mansour L, Andrew Zalesky, Michael Breakspear

## Abstract

The study of functional MRI data is increasingly performed after mapping from volumetric voxels to surface vertices. Processing pipelines commonly used to achieve this mapping produce meshes with uneven vertex spacing, with closer neighbours in sulci compared to gyri. Consequently, correlations between the fMRI time series of neighbouring sulcal vertices are stronger than expected. However, the causes, extent, and impacts of this bias are not well understood or widely appreciated. We explain the origins of these biases, and using *in-silico* models of fMRI data, illustrate how they lead to spurious results. The bias leads to leakage of anatomical cortical folding information into fMRI time series. We show that many common analyses can be affected by this “gyral bias”, including test-retest reliability, fingerprinting, functional parcellations, regional homogeneity, and brain-behaviour associations. Finally, we provide recommendations to avoid or remedy this spatial bias.

## 1. Introduction

Surface-based analysis of fMRI data has several advantages over volumetric analyses, including better correspondence with cortical topology, and the avoidance of artefacts from smoothing across gyral banks (Anticevic et al., 2008). In surface-based analysis, the brain surface is tessellated into a triangular mesh, represented by vertices and faces. Volumetric data are mapped onto the surface through a diffeomorphic projection from grey matter voxels to vertices (Dale et al., 1999; Fischl, 2012; Fischl, Sereno, & Dale, 1999; Fischl, Sereno, Tootell, et al., 1999).

Recent work has found that commonly used template surfaces possess uneven inter-vertex distances, with large variations across different brain regions (Feilong et al., 2023). This problem is not widely appreciated and potentially impacts most surface-based analyses of fMRI data. In the present work, we demonstrate the adverse consequences of this uneven surface projection. First, we show that vertex density is higher in sulci, leading to problems associated with the relative undersampling of non-sulcal regions including gyri. Additionally, because vertex density tracks individuals’ unique cortical folding, structural information can become artefactually incorporated into fMRI, leading to spurious results in commonly used analyses.

It has been previously noted that volume-to-surface mapping of fMRI data results in correlations between neighbouring vertices that are stronger in sulci than in gyri (Ciantar et al., 2022). In particular, when random (spatially uncorrelated) noise time series in volume space are mapped to the surface, adjacent vertices in the sulci have higher correlation than adjacent vertices in the gyri. The bias occurs due to up-sampling from volume data to high-resolution surface meshes. Here we expand on the precise connection between uneven inter-vertex distances (Feilong et al., 2023) and inflated local correlations (Ciantar et al., 2022). We show that inflated local correlations arise not only from volume to surface mapping, but also from current implementations of subsequent surface smoothing.

The gyral bias in diffusion MRI is comparatively well understood. Streamlines in tractography tend to end in gyri rather than sulci (Schilling et al., 2017). In contrast, the consequences of inflated nearest-neighbour correlations in sulci remain poorly understood. We demonstrate that overlooking this problem can lead to spurious results in functional parcellations and fMRI fingerprinting. We recommend remedies to avoid or mitigate this bias.

## 2. Methods

We used minimally preprocessed 3T resting state fMRI data from the Human Connectome Project (HCP), aligned with MSMSulc and cleaned with independent component analysis (Andersson et al., 2003; Andersson & Sotiropoulos, 2015, 2016; Feinberg et al., 2010; Fischl, 2012; Glasser et al., 2013; Glasser & Van Essen, 2011; Jenkinson et al., 2002, 2012; Milchenko & Marcus, 2013; Moeller et al., 2010; Robinson et al., 2014, 2018; Setsompop et al., 2012; Sotiropoulos et al., 2013; Van Essen et al., 2012; Woolrich et al., 2001). Openly available de-identified data from the project were used in accordance with the HCP Data Use Agreement. This study was approved by the University of Newcastle Human Research Ethics Committee (H-2020-0443).

The first 10 participants with 7T fMRI data were used (subject ID 100610, 102311, …). We used one resting state scan (15 minutes) from each participant. Volumetric data were aligned to MNI space. Surface data were represented on the fsLR 32k mid-thickness triangular surface mesh. For parcellation-based analyses, the surface was subdivided into 300 parcels (Schaefer2018_300Parcels_Kong2022_17Networks) (Kong et al., 2021; Schaefer et al., 2018). Sulcal depth values were obtained from the Freesurfer “sulc” output, which measures the height of each vertex above the average surface (Fischl, Sereno, & Dale, 1999).

Vertex pairs directly joined by a mesh edge were termed “neighbours”. Most vertices had 5 neighbours. For each vertex, we calculated the inter-vertex distance as the mean geodesic distance to its neighbouring vertices. We tested the relationship between vertex area and nearest-neighbour correlations. To calculate the area of each vertex’s neighbourhood, we allotted 1/3 of a mesh triangle’s area to each of its 3 vertices. A vertex’s area was calculated as the sum of allotments from all triangles to which it contributed.

Independent random noise time series was used as surrogate data. Standard Gaussian noise time series with 1000 time points were generated independently for each vertex on the fsLR 32k surface. For each vertex, the mean Pearson correlation of its time series with that of neighbouring vertices was termed the “local correlation” of the vertex. This is conceptually similar to regional homogeneity (ReHo), a measure of the rank correlation between the time series of neighbouring vertices (Jiang & Zuo, 2016).

The main results used empirical or random data without additional smoothing. For secondary analyses requiring surface smoothing, we used a 2D Gaussian kernel calculated with geodesic distances between vertices. The kernel was truncated beyond a certain spatial radius, because the kernel is effectively zero for large distances. The fraction of kernel integral discarded by truncation was set to 0.01. This was implemented in the *Connectome Spatial Smoothing* package (Mansour L et al., 2022). Nilearn version 0.10.1 was used for analyses in volume space (Abraham et al., 2014). Pearson’s correlation was used for all associations between two brain maps. Significance was tested using the “spin test”, a spatial permutation test with 1000 permutations implemented in *neuromaps* (Alexander-Bloch et al., 2018; Markello et al., 2022). Volumetric data were projected to a surface using the ribbon-constrained method implemented in Connectome Workbench (Marcus et al., 2011).

## 3. Results

### 3.1. Gyral bias in neighbourhood correlations

Local correlation, the mean time series correlation between a vertex and its mesh neighbours, was highly biased in empirical resting state fMRI data. Sulcal vertices had stronger local correlation than gyral vertices (Figure 1b). The association between local correlation and sulcal depth was significant (r=-0.328, p<0.001).

**Figure 1.**
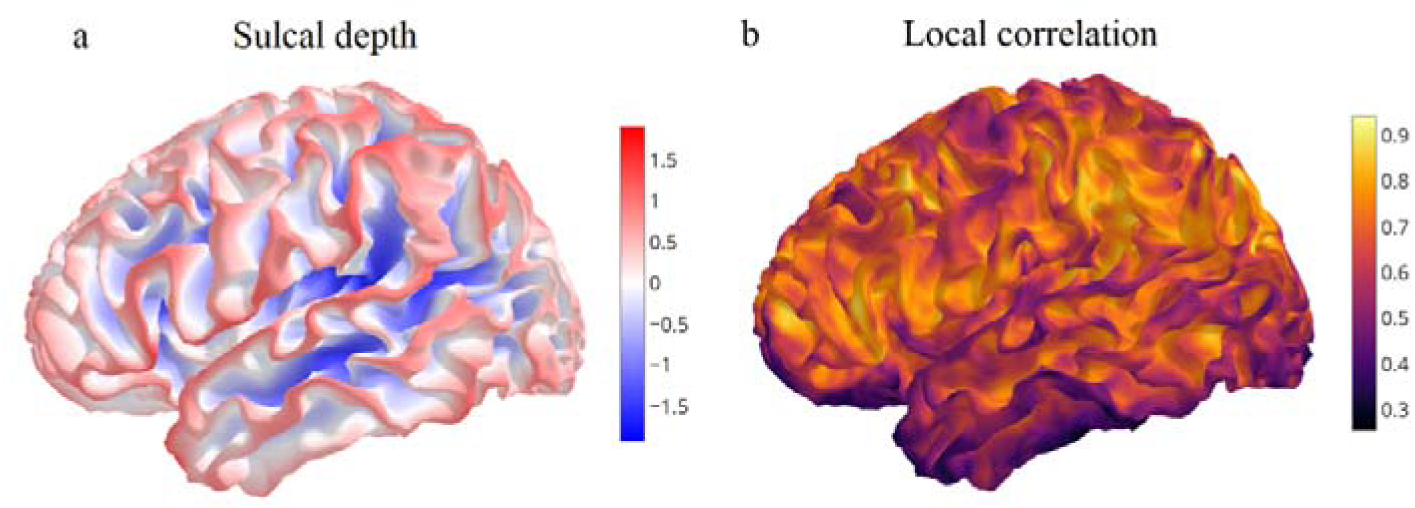
Co-variation of local fMRI correlation with sulci and gyri. Data from a single exemplar participant is shown. Spatial maps are visualised on the participant’s white matter fsLR 32k surface. a) Sulcal depth map with gyri (red) and sulci (blue). b) Resting state fMRI correlation with neighbouring vertices. Correlations were not smoothed, or adjusted for uneven nearest-neighbour distances.

Each vertex’s local correlation value (Figure 1b) was then normalized by subtracting the mean local correlation among all vertices within a 15 mm geodesic radius. This step isolated variation in local correlation values at the fine spatial scale. Normalised local correlation tracked each individual’s gyral crests, following the cortical folding pattern (Figure 2a). Normalised local correlation was associated with sulcal depth (r=-0.500, p<0.001) (Figure 2c). Visualizing an individual’s local correlation map on another participant’s mesh revealed an attenuated association (Figure 2b,d) (r=-0.130, p<0.001). The analysis was repeated with 10 participants. Every participant’s sulcal depth map was more strongly associated with their own local correlation map, than with a different individual’s local correlation map (paired t-test, t(9)=- 19.483, p<0.001) (Figure 2e). Therefore, fMRI neighbourhood correlations contain specific information about an individual’s unique cortical folding. The strong association between local correlations and sulcal depth was replicable with more resting state data (1 hour instead of 15 minutes), MSMAll instead of MSMSulc, and with 7T movie viewing data instead of resting state (Supplementary Figure 1).

**Figure 2.**
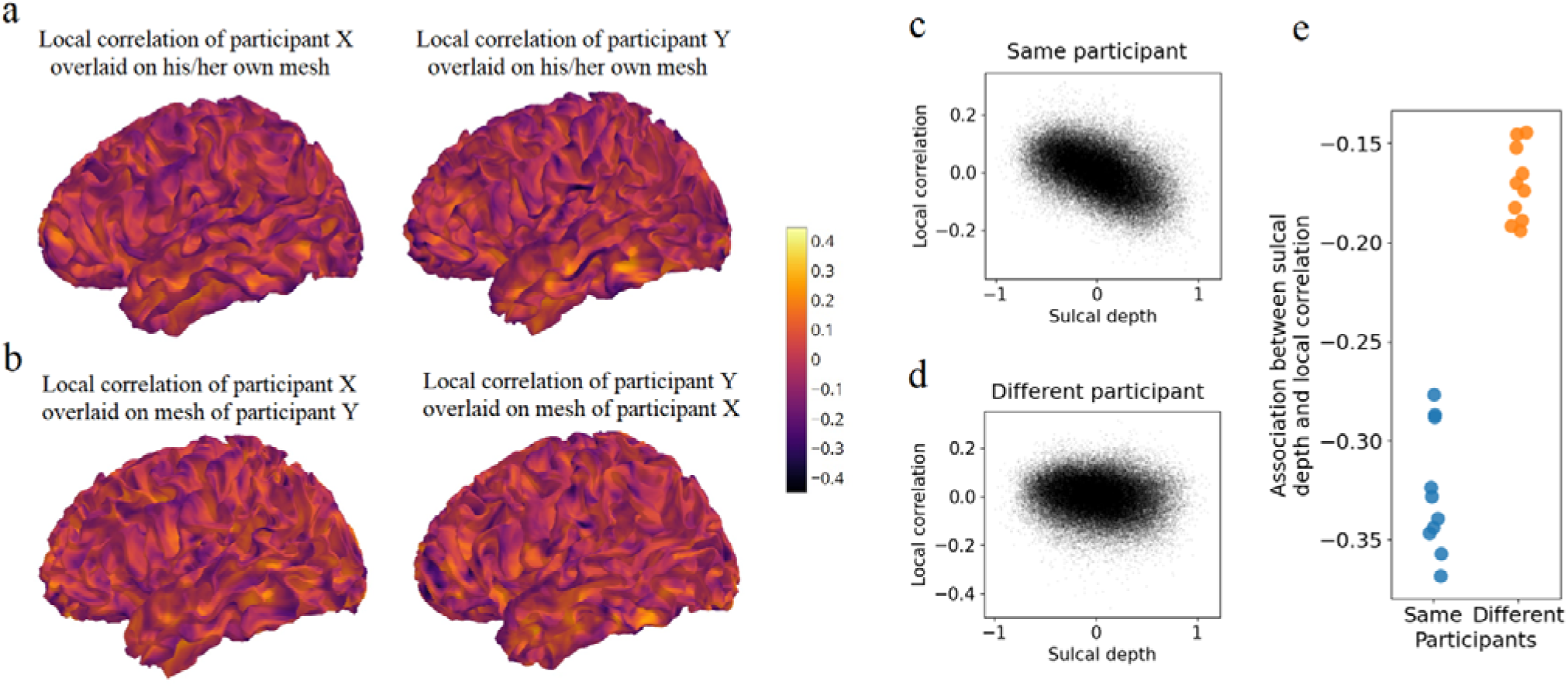
fMRI local correlations track individuals’ unique cortical folding. a) Normalised fMRI local correlation maps for two participants overlaid on their own cortical surfaces. b) Normalised fMRI local correlation maps overlaid on a different participant’s mesh. c) Vertex-wise normalised local correlation values plotted against the same participant’s sulcal depth. Greater sulcal depth values indicate gyri. r=-0.500, p<0.001. d) Normalised local correlation plotted against a different participant’s sulcal depth. r=-0.130, p<0.001. e) Pearson correlation between sulcal depth map and local correlation map, when the maps belonged to the same participants (blue) or different participants (orange).

### 3.2. Uneven vertex spacing in surface meshes

The primary reason for this bias in fMRI local correlations is that surface vertices are spaced further apart in gyri compared to sulci. Uneven spacing of vertices arises because evenly spaced vertices on the inflated sphere are mapped to unevenly spaced vertices on folded surfaces (see Supplementary Results 3 for details). Inter-vertex distances in the mid-thickness surface positively correlated with sulcal depth (r=0.519, p<0.001) (Figure 3b,c). Distances varied from about 1mm at sulci to 3mm at gyral crests. Vertex areas, being proportional to the square of inter-vertex distances, vary by an even greater factor (Figure 3d). Resting state fMRI correlation between a vertex and its immediate neighbours was negatively associated with inter-vertex distance (r=-0.653, p<0.001) (Figure 3e). This appears self-evident, but many common analyses can inadvertently incorporate local correlations unadjusted for inter-vertex distances (see Section 3.3).

**Figure 3.**
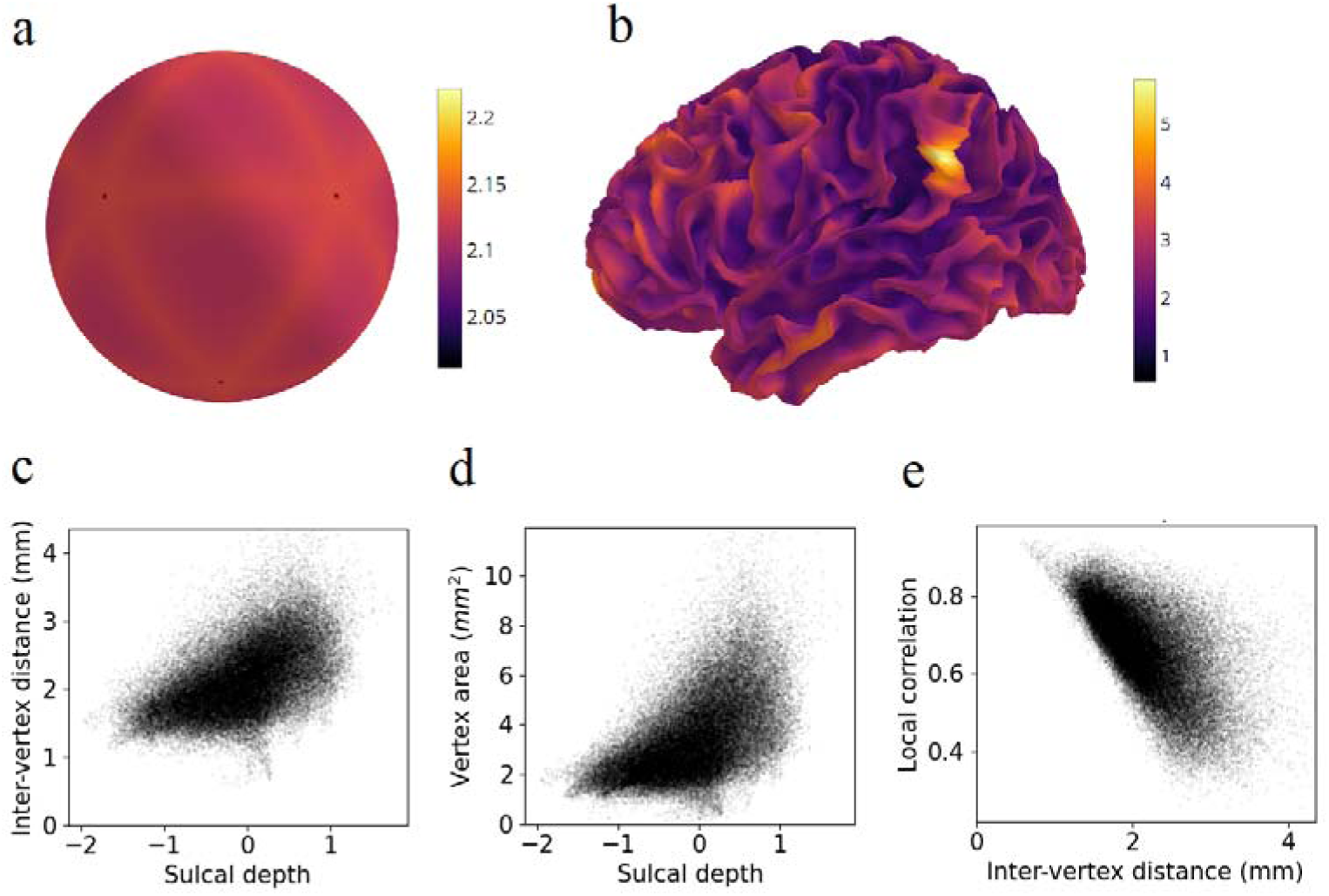
Inter-vertex distance and local correlations: a) Inter-vertex distances on the inflated sphere. b) Inter-vertex distances on the mid-thickness surface. c) Inter-vertex distance on the mid-thickness surface plotted against sulcal depth. Higher sulcal depth values indicate gyri. d) Vertex area plotted against sulcal depth. e) Local fMRI correlation plotted against inter-vertex distance.

There are two mechanisms by which unevenly spaced surface vertices lead to biased fMRI local correlations. These are (i) volume to surface mapping, and (ii) surface smoothing.

#### 3.2.1. Volume to surface mapping

When fMRI data are mapped from MNI space (2mm voxels) to a high-resolution mesh such as the fsLR 32k surface, it is upsampled in some areas. In regions with low inter-vertex distances such as sulci, one voxel is upsampled to more than one vertex. Thus, adjacent vertices will have the same or similar time series (Ciantar et al., 2022). This does not occur in regions with high inter-vertex distances such as gyri. To simulate this, we generated uncorrelated fMRI noise time series in MNI volume space, independently for each voxel. These time series were projected to a single participant’s mid-thickness surface. The resulting surface time series had greater local correlation within sulci (Figure 4b). Vertices with higher inter-vertex distance had lower local correlations (r=-0.596, p<0.001) (Figure 4c).

**Figure 4.**
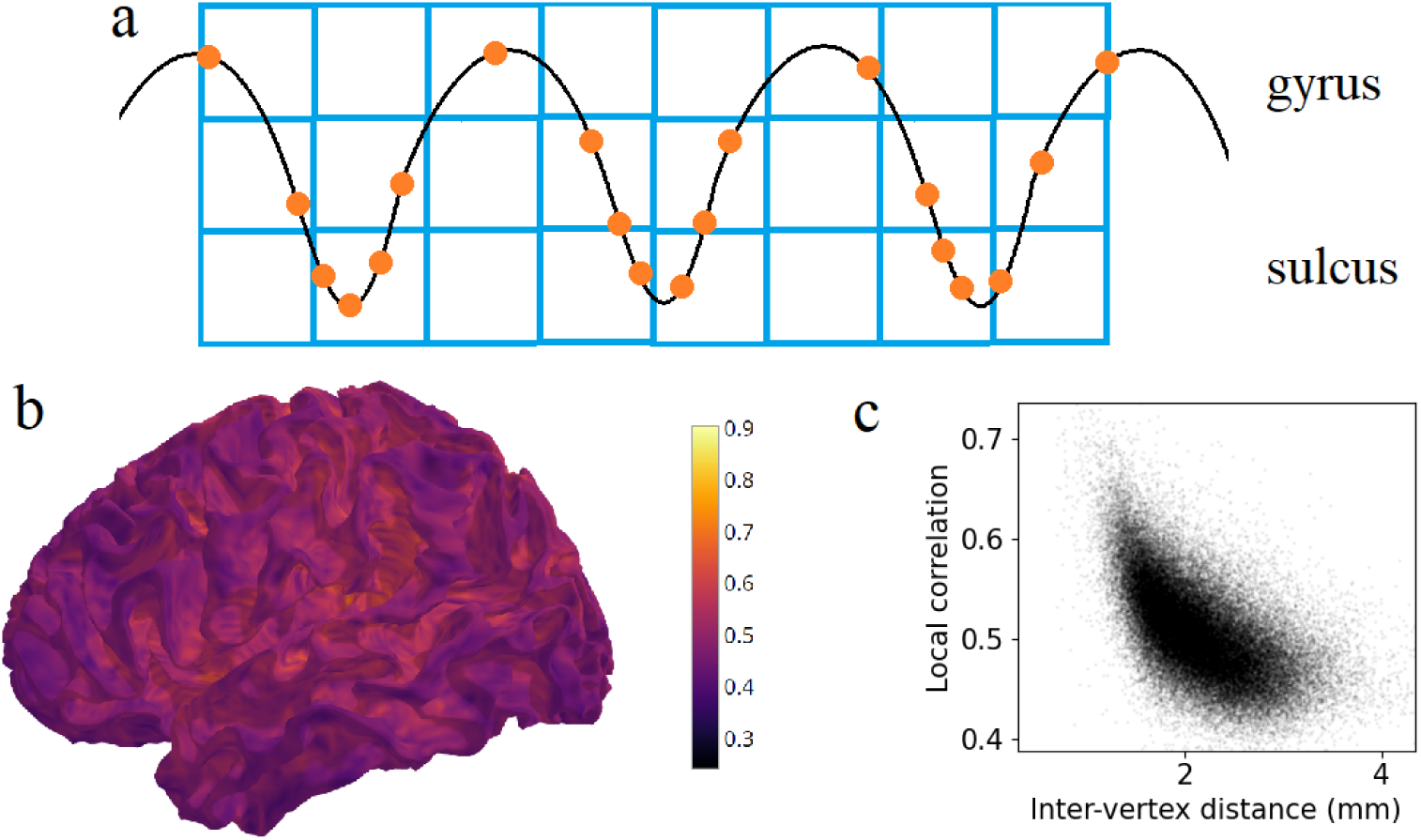
Volume to surface mapping induces artefactual local correlations. a) Schematic showing that a single voxel’s (blue square) data is upsampled to multiple vertices (orange dot) due to small inter-vertex distances, especially in sulci. b) Uncorrelated fMRI noise time series in volume MNI space were projected to the mid-thickness surface. This plot shows local correlations between adjacent vertices in the surface space. c) Correlation between adjacent mesh vertices against inter-vertex distance.

#### 3.2.2. Surface smoothing

Surface smoothing independently induces artefactual local correlations. Since vertices are closer to one another in sulci than in gyri, smoothing has greater effect on local correlations in sulci (Figure 5a). To illustrate this, we generated uncorrelated noise time series, this time in surface space, independently for each vertex. These time series were smoothed on the surface with a 2mm FWHM kernel. Surface fMRI data is often smoothed at this level (Glasser et al., 2013). This resulted in greater local correlation within sulci than in gyri (Figure 5b), and a negative association between inter-vertex distance and local correlations (r=-0.867, p<0.001) (Figure 5c). There was no such bias when smoothing was applied to white noise generated on the spherical surface, since vertices are approximately evenly spaced on the sphere (r=0.007, p=0.208).

**Figure 5.**
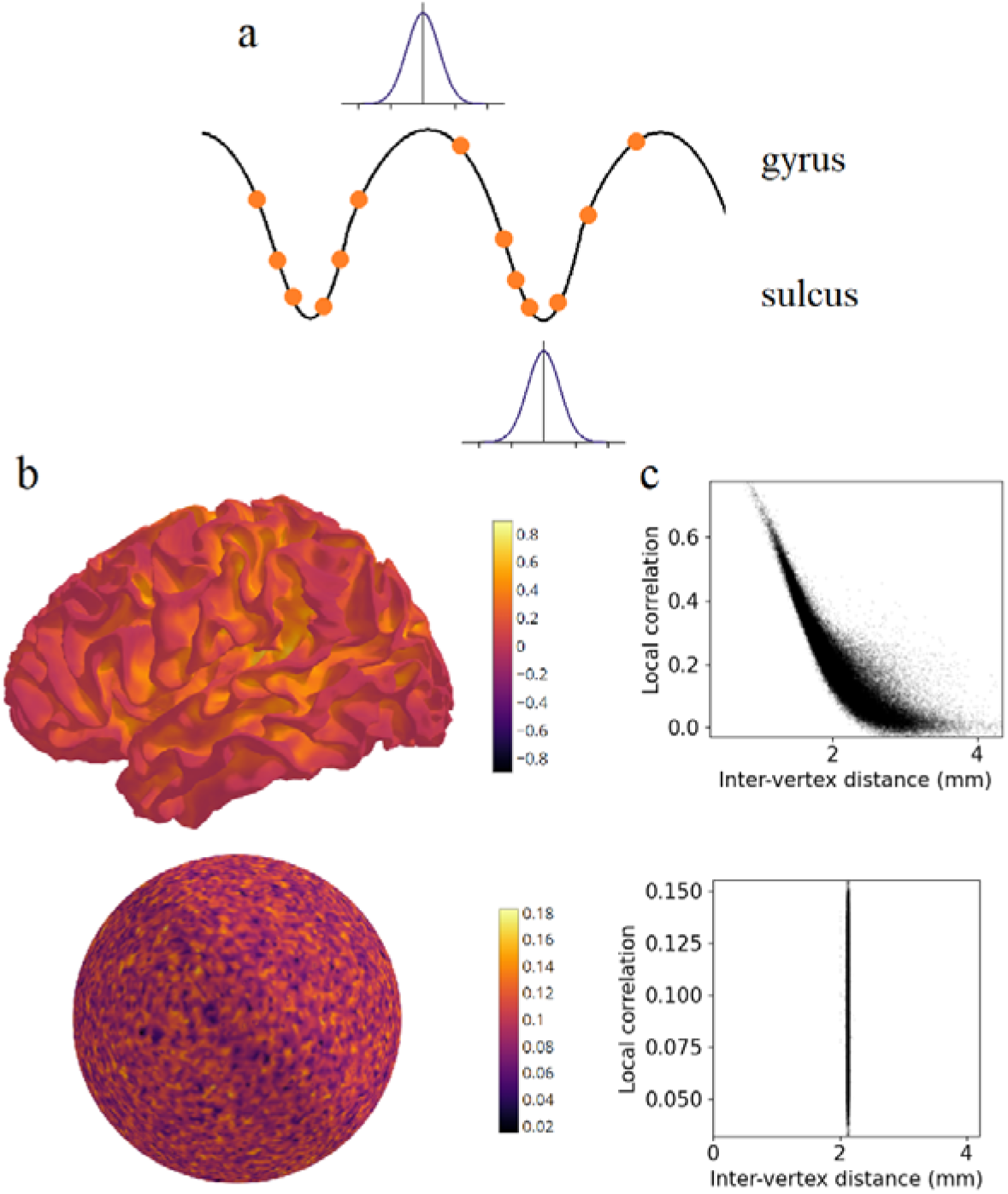
Surface smoothing induces artefactual local correlations. a) Schematic showing that a Gaussian kernel of fixed width smooths over multiple vertices in sulcal regions due to small inter-vertex distances. b) Uncorrelated noise time series generated on the mid-thickness surface (top) or spherical surface (bottom) were smoothed with a 2mm FWHM Gaussian kernel. This plot shows local correlations between adjacent vertices in the surface space. c) Correlation between adjacent vertices against inter-vertex distance, on the mid-thickness surface (top) or spherical surface (bottom).

The artefactual correlations induced by volume to surface mapping can be avoided by using a lower resolution surface mesh (Ciantar et al., 2022). However, surface smoothing induces spurious local spatial structure even when a lower resolution mesh is used. In the group-averaged standard fsaverage5 pial surface, surface smoothing of noise data resulted in biased local correlations associated with inter-vertex distances (r=-0.726, p<0.001) (Supplementary Figure 2).

#### 3.2.3. Gyral bias in surface data does not originate from biases in volume data

It is plausible that differences in local correlations between gyri and sulci arise not from surface processing, but from earlier preprocessing steps, or from ground truth effects such as blood flow differences. We therefore tested for biases in empirical HCP resting state fMRI volumetric data, where voxels are spaced uniformly in MNI space. For each voxel, mean correlation with its 6 neighbouring vertices (anterior, posterior, superior, inferior, left, right) were calculated. Vertices outside the grey matter were excluded. The map of local correlation values was calculated in volume space, then projected to the participant’s mid-thickness surface for visualisation. Local correlations in volumetric space were not significantly associated with sulcal depth (r=0.012, p=0.762) (Supplementary Figure 3). This result suggests that the gyral bias in surface data arises from data processing rather than being a feature of the BOLD response.

We next queried whether gyral bias on surface-mapped fMRI would persist if more distant vertices were considered, and if the data were adjusted for inter-vertex distances. To this end, empirical resting state fMRI data was used to estimate, for each vertex, spatial autocorrelation as a function of distance. Spatial autocorrelation was not greater in sulci compared to gyri when adjusted for distance, rather than calculated amongst local vertices (Supplementary Results 4).

### 3.3. Consequences of gyral bias in functional fMRI

In the previous section, we demonstrated how uneven vertex spacing in the discretised cortical surface leads to artefactual correlations. In this section, we provide several examples of how fMRI gyral bias can contaminate surface-based neuroimaging analyses.

#### 3.3.1. Surface parcellations based on functional MRI

Functional parcellation schemes aim to divide the brain into contiguous parcels with relatively homogenous fMRI signals within each parcel. We hypothesized that the borders of a surface functional parcellation would be biased to prefer gyri over sulci. Parcel boundaries prefer areas where fMRI signal changes rapidly (gyri), while parcel centres tend to be located at vertices that are similar to many surrounding vertices (sulci).

To illustrate this, we constructed a null dataset for a single participant by generating random noise time series for each vertex on the participant’s fsLR 32k surface. The time series had 500 time points. The data (D^59412^ ^x^ ^500^) were smoothed by a 2mm FWHM Gaussian kernel. The preceding analysis in Section 3.2.2 demonstrated how this yields high local correlations in sulci. Thus, the correlations in the data implicitly contained participant-specific cortical folding information, rather than true functional information.

We used a single participant’s null data constructed in this manner. Ward’s hierarchical agglomerative clustering algorithm (Johnson, 1967) was used to find 50 clusters in the left hemisphere (Thirion et al., 2014). After vertices were assigned to parcels, we identified border vertices as those which had a neighbour in a different parcel. Border vertices have a more positive mean sulcal depth value (0.145) compared to non-border vertices (-0.159), indicating that they are closer to gyri on average. The difference in sulcal depth values between border and non-border vertices was confirmed with a two-sample unpaired t-test (Cohen’s d=0.540, t(29694)=41.636, p<0.001 with spin test) (Figure 6). We then repeated the analysis in surrogate data generated on the surface of 10 other participants. All participants had a t-statistic greater than 20, indicating that border vertices were closer to gyri. A second-level one-sample t-test was employed to test whether the 10 participant-specific t-statistics deviated from zero. The effect was significant (t(10)=20.164, p<0.001), indicating that border vertices systematically align with gyri across participants.

**Figure 6.**
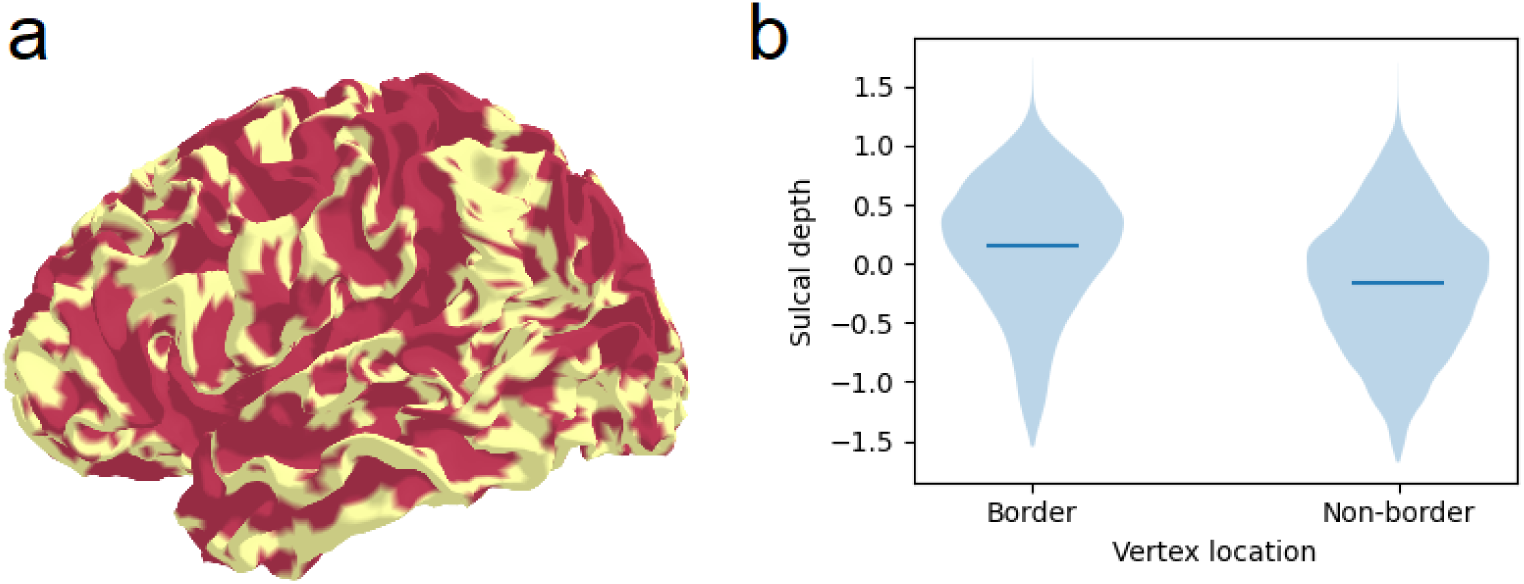
Boundaries of a functional parcellation are closer to gyri than to sulci. a) (left) Border vertices (pink) and non-border vertices (yellow) in one participant’s parcellation. b) (right) Sulcal depth values in border and non-border vertices. Positive sulcal depth values indicate gyri. Horizontal lines indicate mean.

We hypothesized that subject-specific cortical folding can contribute to the “individuality” in individualized functional parcellations. This is not necessarily the case, for example if functional parcel borders follow smoother group-averaged patterns of cortical folding but not individuals’ unique gyrifications. Following our hypothesis, we predicted border vertices calculated from one participant would not align with the gyri of a different participant. We tested the correspondence between the parcel borders of the 10 participants and the sulcal depth maps of 10 *different* participants. This yielded a new set of 10 t-statistics. The new “mismatched” set of t-statistics were smaller than the original “matched” set (paired t-test, t(9)=8.121, p<0.001), suggesting that individualised functional parcellations are influenced by the individual cortical folding information embedded in the local correlations of fMRI data.

We next tested whether these results held in commonly used group functional parcellations derived from empirical data. Parcel borders were closer to gyri in the Schaefer 300-node parcellation. Border vertices had greater sulcal depth values than non-border vertices (Cohen’s d=0.127, t(29694)=9.801, p<0.001 with spin test). The effect size was greater with the 100-node parcellation (Cohen’s d=0.218, t(29694)=14.306, p<0.001 with spin test). We then tested whether cortical folding influences the individuality of parcel boundaries in empirical individualized parcellations. In a commonly used 300-node individualized parcellation (Kong et al., 2021), an individual’s parcel boundaries followed the same individual’s cortical gyri more closely than the cortical gyri of a different individual (paired t-test, t(9)=4.345, p<0.001).

#### 3.3.2. Test-retest reliability and fingerprinting in functional connectivity

“Fingerprinting” in fMRI refer to the ability to identify a given individual’s follow-up fMRI scan from within a large set of other participants’ scans, if one is provided the same individual’s baseline scan. Fingerprinting relies on low within-subjects variability (or high test-retest reliability) and high between-subjects variability.

It is known that cortical folding patterns are highly reliable within subjects and differ sufficiently between participants to allow fingerprinting (Pizzagalli et al., 2020). We have shown how local correlations track an individual’s cortical gyrifications. Therefore, vertex-resolution functional connectivity could potentially distinguish individuals by virtue of having structural information embedded in fine-scale correlation patterns.

As in section 3.3.1, we used null data generated on the surface meshes of 10 different participants, constructed by generating spatially independent noise time series on the surface and smoothing it with a 2mm FWHM kernel. Two such “scans” were generated for each participant, corresponding to test and re-test scans. Using these data, we investigated the fingerprinting accuracy of parcel-level and vertex-level functional connectivity separately. For the parcel level, we averaged across vertices within each parcel to obtain the parcels’ time series (D ^300^ ^x^ ^500^). The parcel-level functional connectivity matrix (F ^300^ ^x^ ^300^) was calculated by taking the Pearson correlation between parcels’ time series. For the vertex level, we only considered vertices within a single parcel. This was because computing the full 59,412 x 59,412 vertex-level connectome would be computationally demanding, and unnecessary for our demonstration. We therefore computed functional connectivity within a single parcel (parcel 1) in the 300-node parcellation. The time series for these vertices was represented by D_V_^N^ ^x^ ^500^, where N was the number of vertices within that parcel. The vertex-level functional connectivity matrix (F_V_^N^ ^x^ ^N^) was calculated using the Pearson correlation between vertex’s time series. Functional connectivity matrices were vectorised.

Given a pair of participants A and B, we calculated the Pearson correlation between participant A’s test functional connectivity matrix, and participant B’s re-test functional connectivity matrix. This was repeated for all choices of participants A and B. Since test and re-test scans were generated as independent noise, the test-retest correlation should be similar irrespective of whether A=B or A≠B. This is the case for parcel-level functional connectivity. However, at the vertex level, each participant’s re-test scan was more strongly correlated with their own test scan than other participants’ test scans. We then sought to quantify fingerprinting accuracy. Each participant’s retest scan was matched with the test scan with which it was most highly correlated. Fingerprinting accuracy was quantified by the proportion of retest scans which were correctly matched. Accuracy was 10% (1/10) at the parcel level, but 100% (10/10) at the vertex level despite test and retest scans being uncorrelated noise (Figure 7a).

**Figure 7.**
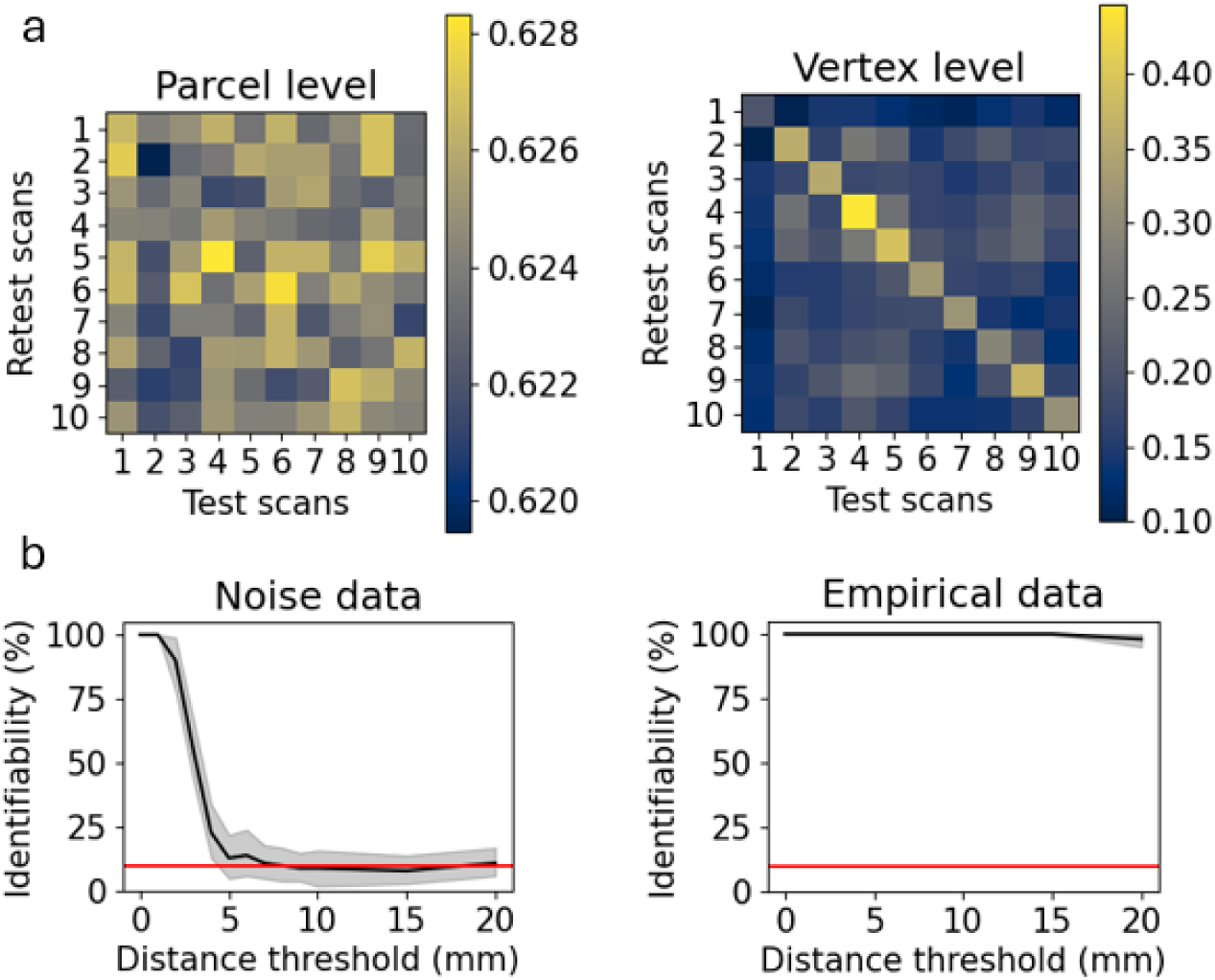
a) Fingerprinting accuracy of parcel-level (left) and vertex-level (right) functional connectivity in smoothed random noise data. Colours indicate the correlation coefficient between a participant’s test scan and all participants’ retest scan. Correlations were calculated between vectorised functional connectivity matrices. In each plot, the horizontal axis indicates which participant’s test scan was used. The vertical axis indicates which participant’s retest scan was used. b) Fingerprinting accuracy of vertex-level functional connectivity in single parcels as a function of distance threshold, for smoothed random noise (left) or empirical resting state fMRI (right). Functional connectivity data was omitted for vertices closer than the distance threshold. Fingerprinting accuracy values were averaged across parcels. Shading indicates 95% confidence interval. Horizontal line indicates chance level (10%).

We tested whether vertex level fingerprinting accuracy depends on correlations between nearby vertices. Short-range connections were removed from the functional connectivity matrix. A threshold value for geodesic distance was selected. Elements of matrix F_V_^N^ ^x^ ^N^ were removed if they corresponded to vertex pairs closer than the distance threshold. Fingerprinting accuracy dropped to chance level when the distance threshold was above 5mm. In empirical resting state data, identifiability remained high beyond the 5mm distance threshold (Figure 7b).

Considering this result, studies examining test-retest reliability and fingerprinting of fine-scale functional connectivity should be cautious of structural information embedded in fine-scale fMRI, especially when including correlations between nearby vertices. Heritability of fine-scale functional connectivity is another area of concern, as cortical folding patterns are heritable (Pizzagalli et al., 2020). Notably, fingerprinting of parcel-level functional connectivity derived from smoothed random noise is no higher than chance. However, parcel-level data may still convey structural information. For example, if a sulcus is located near a border between two parcels, then vertices on opposite sides of the border will be correlated. This increases the correlation between the two parcels. In this way, sulcal location information can remain embedded in the functional connectivity between adjacent parcels.

#### 3.3.3. Sulcal vertices contribute disproportionately to parcels

Parcel-level analyses rely on averaging the values of all vertices within a parcel. We have shown that inter-vertex distances in gyri are 2-3 times greater than in sulci. This means that sulcal vertices are over-represented in surface space. Therefore, parcel mean values will under-represent gyral data. This issue is relevant to analyses that employ vertex-level data including fMRI, vertex-wise genetic maps, or spatial maps of myelin thickness. As an illustration, we generated a single participant’s null data as in section 3.3.1 by surface smoothing a spatially independent white noise time series. Parcel mean time series were calculated for each parcel in the Schaefer 300-node parcellation. For each vertex, we calculated the Pearson correlation between its time series and the time series of its encompassing parcel. This reflects the contribution of each vertex to its supervening parcel. We tested whether a vertex’s sulcal depth was associated with its correlation with the parcel mean. Gyral vertices in this test data had significantly lower correlations with parcel means compared to sulcal vertices (r=-0.249, p<0.001), indicating that they are under-represented at the parcel level (Supplementary Figure 5). Parcel mean time series were then re-calculated using a weighted average, where each vertex’s value was multiplied by the estimated vertex area. This procedure reduced the magnitude of the bias but did not abolish it (r=0.102, p<0.001).

Biased parcel means occurred for the following reason: Uncorrelated noise time series in sulcal vertices became correlated after surface smoothing. Gyral vertices remained relatively uncorrelated because they are distant relative to the 2mm FWHM smoothing kernel. When parcel means are calculated, the time series of uncorrelated gyral vertices tend to cancel out. Conversely, the time series of correlated sulcal vertices contribute collectively to the parcel average. This example demonstrates that sulcal vertices can contribute disproportionately to surface-based parcel level analyses.

## Discussion

We studied the influence of heterogenous vertex spacing in commonly used surface meshes which possess greater inter-vertex distances in gyri compared to sulci. This gyral bias in inter-vertex distances yields a gyral bias in nearest neighbour correlations. In particular, uneven vertex spacing and local correlations track not only group-averaged cortical folding, but also individuals’ gyrifications. Consequently, individual-specific structural information becomes embedded in the corresponding individual fMRI. Using simulated noise, we demonstrated how spurious results can arise in the case of functional parcellations, fingerprinting, and parcel averages.

In many commonly used surface spaces, vertices are positioned more densely in sulci, with sulci being oversampled by a factor of 3 or more relative to gyri. This stems from preprocessing steps that have been standardised across the field (Supplementary Results 1). Consequently, surface-based functional parcellations can be susceptible to fMRI gyral bias. Most functional parcellations directly cluster vertex time series, use fMRI connectivity gradients to find parcel boundaries, or combine both approaches. Direct clustering leads to parcel centroids that are biased towards sulci, because sulcal data is over-represented. fMRI connectivity gradients, when uncorrected for inter-vertex distances, rapidly change in gyri due to low local correlations, leading to parcel boundaries biased towards gyral loci. In vertex level functional connectivity, we found that smoothed noise time series are spuriously identifiable if correlations between nearby vertices (within 5mm) are included. We explain the 5mm threshold by noting that it is approximately equal to the largest inter-vertex distances on the fsLR 32k cortical mesh (Figure 3). Some gyral vertices are 5mm from their nearest neighbours, while sulcal vertices have much closer nearest neighbours. Omitting short-range connections removes this distinguishing feature of sulcal vertices, and reduces the influence of cortical folding patterns on fMRI correlations. In general, we advise researchers against calculating fMRI correlations between vertices that are within a critical distance threshold, the largest inter-vertex geodesic distance. This is about 5mm for the fsLR 32k mesh.

There are several other scenarios where analyses can be confounded by these biases. These fall into two categories - situations arising from oversampling of sulci and those arising from embedding structural information in fMRI correlations.

There are several other possible consequences of oversampling sulci. First, relative under-sampling of gyri could impact predictive accuracy in multi-voxel pattern analysis or brain-behaviour prediction. Second, it can lead to problems with hyperalignment (Haxby et al., 2011, 2020) (also known as functional alignment). This method estimates a transformation matrix that maps vertices in one participant to vertices in another. The optimal matrix transforms the first participant’s functional response to predict the second participant’s response. There can be constraints on the matrix’s form. For instance, a permutation matrix maps single vertices to single vertices. In optimal transport, each vertex contributes equal “mass” which is divided among target vertices in the output space (Bazeille et al., 2019). These approaches assume the equivalence of gyral and sulcal vertices. The assumption is not substantiated in surface meshes, where a gyral vertex typically occupies a surface area that is the equivalent of multiple sulcal vertices.

Confounding of results can also arise from structural information embedded in the local correlations of fMRI. Any associations between fMRI and other data modalities could be influenced by gyral biases being common to both. While we have shown artefactual gyral biases in surface fMRI data, similar biases are present in other modalities. Cortical thickness is greater in gyri (Fischl & Dale, 2000). This is a real anatomical difference. Diffusion MRI contains an artefactual gyral bias wherein streamlines tend to end at gyri (Schilling et al., 2017). EEG and MEG are more sensitive to gyral source and sulcal walls respectively due to different orientations of the dipole fields (Srinivasan, 2006). Testing for associations between these different modalities must be approached with caution. Regional homogeneity (Jiang & Zuo, 2016), a measure of fMRI correlation between neighbouring vertices, would also be artefactually elevated in sulci when calculated on surface fMRI data. The distribution of fMRI correlations across the cortex could be influenced by several other factors, including regional differences in fMRI signal-to-noise ratio (Jezzard & Clare, 1999), head motion, and the distribution of vasculature (Huck et al., 2023).

Finally, when predictive models are trained on vertex-resolution fMRI to predict behaviour or cognitive phenotypes, implicit structural information may be contributing to the prediction. For instance, a neural network may learn to use the correlation between adjacent vertices (which tracks cortical folding) as a feature for prediction.

There are several approaches to excluding fMRI gyral biases. First, using a surface mesh with resolution less than that of the voxel data avoids the problem of upsampling during volume-to-surface mapping (Ciantar et al., 2022). For example, projecting from 2mm MNI voxel space to the coarse fsaverage5 surface (with 10k vertices) avoids projecting one voxel onto multiple vertices. However, even at these lower resolutions, surface smoothing will induce artefactual local correlations. Another possibility is to conduct analyses such as parcellation in volume space before projecting results to the surface (Ciantar et al., 2022). However, this eschews the benefits of surface representations. For instance, a volume-based parcellation could misconstrue voxels on opposite sulcal banks as adjacent. A recent promising approach has been the introduction of surface templates with more homogenous inter-vertex distances. The *onavg* template, developed through an iterative procedure of adjusting vertex locations, reduces inter-vertex spacing by about 90% compared to standard meshes (Feilong et al., 2023). This template samples gyri and sulci more uniformly and improves predictive accuracy in multi-voxel pattern analysis. However, some residual variability in inter-vertex distance remains. A “perfect” group-level template is impossible because vertex locations with uniform inter-vertex distances in one participant would not have the same uniform spacings in another. For within-participant analyses like multi-voxel pattern analysis, generating participant-specific meshes using the same iterative method (Feilong et al., 2023) may be the best option.

Excluding the fMRI gyral bias is an ongoing challenge. Researchers should use quantitative measures that are robust against uneven vertex spacing. For instance, spatial autocorrelation as a function of geodesic inter-vertex distance is robust, while nearest-neighbour correlation is not robust to this bias. Computing parcel means with a weighted average proportional to vertex area attenuates the bias. Even with these modifications, the influence of fMRI gyral biases may remain obscure in complex multi-step pipelines (Supplementary Results 2). In this situation, researchers can still test whether fMRI gyral bias can explain study results. By running a pipeline on gyrally biased random noise data instead of empirical data, one can test whether the observed finding arises spuriously from randomly generated null models. Surface-smoothed Gaussian noise data as studied here can be used for this purpose. Finally, any study results should be visualised on participant-specific non-inflated surfaces, so that obvious covariation with sulcal depth can be identified.

Cortical data benefit greatly from surface representations due to the cortex’s folded sheet-like topology. However, researchers should remember that cortical meshes use a somewhat arbitrary tessellation to discretise the brain surface. Care should be taken that analyses do not implicitly treat vertices as ground truth entities or features.

## Data and code availability

Code to perform analyses is available at https://github.com/jaysonjeg/track_align. Data were from the Human Connectome Project and can be accessed at https://db.humanconnectome.org/

## Author contributions

JJ – conceptualization, methodology, software, analysis, visualization, original draft, review and editing. BP and NK – software, original draft, review and editing. SML and AZ – methodology, review and editing. MB – conceptualization, methodology, original draft, review and editing.

## Acknowledgements

Data were provided [in part] by the Human Connectome Project, WU-Minn Consortium (Principal Investigators: David Van Essen and Kamil Ugurbil; 1U54MH091657) funded by the 16 NIH Institutes and Centers that support the NIH Blueprint for Neuroscience Research; and by the McDonnell Center for Systems Neuroscience at Washington University.

