## Supplementary Figure 1 for "Spurious correlations in surface-based functional brain imaging"

**Supplementary Materials**

Supplementary Results 3. FreeSurfer and fsLR spherical registration and downstream gyral bias

The FreeSurfer spherical registration method offers several advantages over volumetric registration, with a robust method for inter-subject registration of fMRI studies. After cortical reconstruction of pial and white surfaces, these are inflated using an energy-minimization algorithm (Fischl et al., 1999) that minimizes metric distortions of the surfaces. These participant-specific spheres are then registered to the FreeSurfer spherical template space, allowing alignment to a common space for intersubject comparison. However, the transformation of cortical surfaces to this common space induces a gyral bias. Vertices are nearly uniformly spaced on the FreeSurfer template sphere, but this corresponds to unevenly spaced vertices in the subsequent folded cortical surfaces. This arises from the method of “re-folding” the cortex from the sphere.

As an example, we highlight the steps in the HCP Minimal Preprocessing Pipeline that are involved in these transformations (Glasser et al., 2013). The pipeline distinguishes a *Native* coordinate space that is specific to the subject’s cortical reconstructions; these reconstructions being derived from FreeSurfer prior to FreeSurfer spherical registration (i.e., using *recon-all* up to --*autorecon3*). These reconstructions have unevenly spaced vertices but express no discernible pattern to this spacing). In the unmodified *HCPpipelines* post-FreeSurfer registration step contained here: https://github.com/Washington-University/HCPpipelines/blob/master/PostFreeSurfer/scripts/FreeSurfer2CaretConvertAndRegisterNonlinear.sh, with default parameters, *Native* surfaces are saved to the filepath “{Subject}/T1w/Native”. Registration of these surfaces to the *fsLR* template then proceeds to a (i) high-resolution *164k* space with 163,842 vertices per hemisphere, equivalent to the *fsaverage* space (also with 163,842 vertices; however, these were reordered in *fsLR* to be roughly equivalent across hemispheres (Van Essen et al., 2012), and to a (ii) low-resolution *32k* space, with 32,492 vertices per hemisphere.

Both template spheres, *164k* and *32k*, have nearly uniform inter-vertex spacing. The (i) registration to the fsLR *164k* template utilises the FreeSurfer template-registered individual subject spheres (copied from the individual’s FreeSurfer “{l/r}h.sphere.reg” to “{Subject}/MNINonLinear/{Subject}.{Hemisphere}.sphere.164k_fs_LR.surf.gii”). As these spheres are registered to the FreeSurfer template space, the gyral bias makes its way to the subject’s “MNINonLinear” surfaces. These surfaces are contained in the filepath “{Subject}/MNINonLinear/”, “{Subject}/MNINonLinear/Native”.

Registration in (ii) proceeds by *Native* registration to the *32k* downsampled template sphere. The definition of vertex points in the *32k* template sphere was obtained similarly to the *fsaverage* template spheres. The template sphere was parameterized as a geodesic polyhedron (in this case, a subdivided icosahedron with a spacing of 2.02-2.18 mm; see Figure 3a). In the HCP Pipelines (with default parameters) this template sphere is contained in https://github.com/Washington-University/HCPpipelines/blob/master/global/templates/standard_mesh_atlases/L.sphere.32k_fs_LR.surf.gii. This file is copied to “{Subject}/MNINonLinear/fsaverage_LR32k/{Subject}.{Hemisphere}.sphere.32k_fs_LR.surf.gii” during the *PostFreeSurfer* pipeline. While the points are evenly spaced on the spherical surface, the sampling of points in the resultant *32k* folded cortical surfaces is far denser in sulci than gyri.

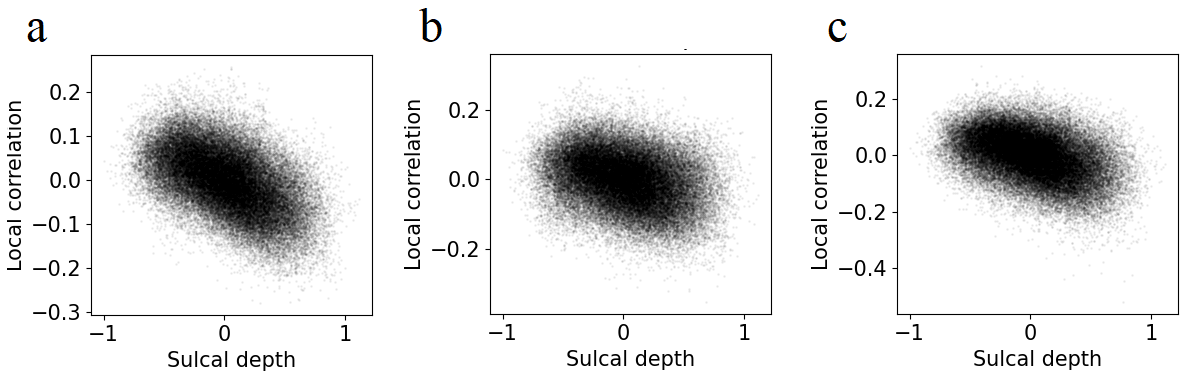

Supplementary Figure 1. fMRI local correlations plotted against sulcal depth. Higher sulcal depth values indicate gyri. Both variables were normalised by parcel-specific mean values. fMRI local correlations tracked individuals’ unique cortical folding, irrespective of changes to the fMRI data type. a) Using 4 runs of resting state fMRI (1 hour) instead of 1 run (15 minutes) (r=-0.525, p<0.001). b) Using resting state data aligned with MSMAll instead of MSMSulc (r=-0.294, p<0.001). c) Using 7T movie viewing fMRI (1 run) (r=-0.413, p<0.001).

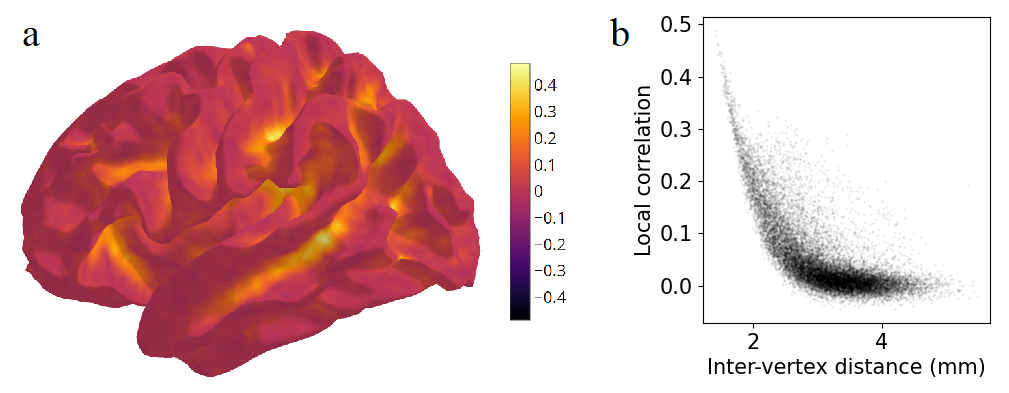

Supplementary Figure 2. Surface smoothing (2mm FWHM) induces biased fMRI correlations in the fsaverage5 pial surface. Uncorrelated noise was generated at each surface vertex. a) Local correlation at each vertex. b) Local correlation plotted against inter-vertex distance (r=-0.726, p<0.001).

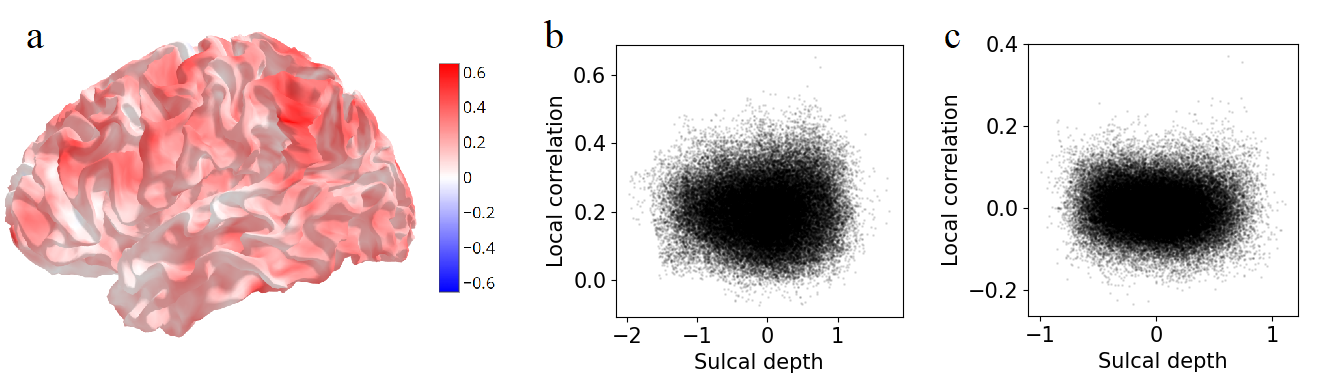

Supplementary Figure 3. Local correlations in resting state fMRI in volume space. Volume-based local correlation were projected to the surface for visualisation. a) Local correlations. b) Local correlation plotted against sulcal depth (r=0.012, p=0.762). Higher sulcal depth values indicate gyri.

Supplementary Results 4. Spatial autocorrelation as a function of distance

We used empirical HCP resting state fMRI data in the fsLR 32k surface space. The following pipeline was followed to estimate a different spatial autocorrelation function for each vertex. The original vertex was termed the “source vertex”. We considered an expanded definition of vertex neighbourhood, and all other vertices within a 10mm geodesic distance were termed “neighbours”. We calculated the fMRI correlation between the source vertex time series and its neighbours’ time series, and the geodesic correlation between them on the mid-thickness mesh. These data characterized the source vertex’s spatial autocorrelation function. An exponential curve $y=Ae^{-kx}+c$ was fitted to predict correlation from distance. Parameter $k$, the decay rate, indicates how rapidly spatial autocorrelation drops with distance from the source vertex. We calculated the decay rate for each vertex on the fsLR 32k surface. Decay rate is reduced in gyri compared to sulci (r=-0.191, p<0.001), indicating that the spatial extent of spatial autocorrelation is larger in gyri (Supplementary Figure 4).

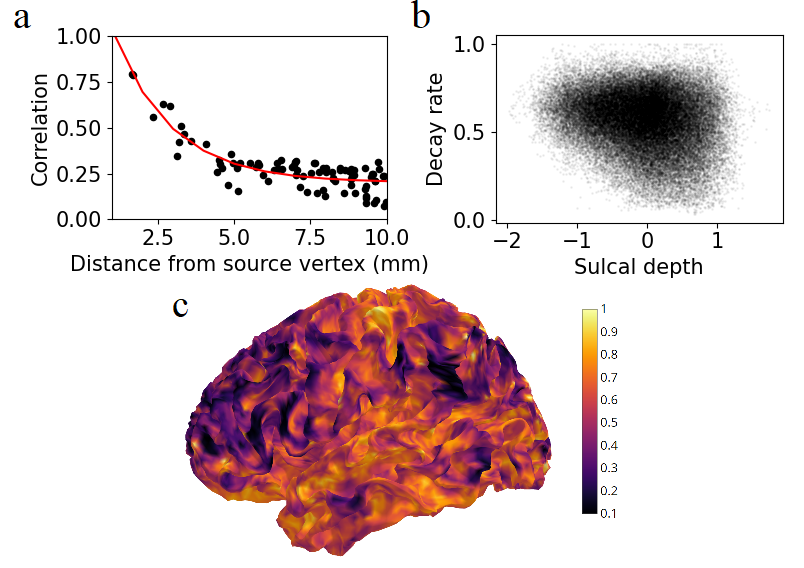

Supplementary Figure 4. Spatial autocorrelation as a function of distance on the mid-thickness surface. a) Example of the spatial autocorrelation function of a single vertex. The red line indicates the fitted exponential function b) The exponential function decay rate is plotted against sulcal depth. Each data point indicates a single vertex. Positive sulcal depth values indicate gyri. d) Spatial map of the decay rate.

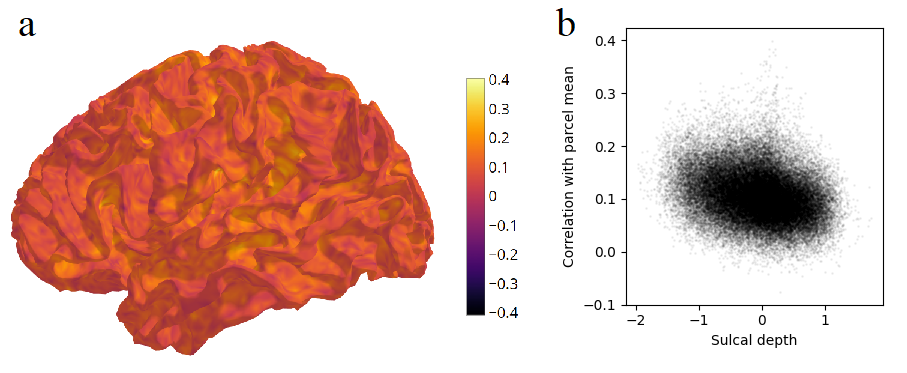

Supplementary Figure 5. Sulcal vertices contribute disproportionately to parcel means. Noise time series were smoothed with a 2mm FWHM Gaussian kernel. a) Correlation between each vertex’s time series and its parcel mean time series. b) Plot of the correlations in (a) against sulcal depth. Positive values indicate gyri. r=-0.249, p<0.001
